# Natural and after colon washing fecal samples: the two sides of the coin for investigating the human gut microbiome

**DOI:** 10.1101/2021.06.29.450302

**Authors:** Elisabetta Piancone, Bruno Fosso, Mariangela De Robertis, Elisabetta Notario, Annarita Oranger, Caterina Manzari, Marinella Marzano, Silvia Bruno, Anna Maria D’Erchia, Dominga Maio, Martina Minelli, Ilaria Vergallo, Mauro Minelli, Graziano Pesole

## Abstract

To date there are several studies focusing on the importance of gut microbiome for human health, however the selection of a universal sampling matrix representative of the microbial biodiversity associated to the gastrointestinal (GI) tract, still represents a challenge. Here we present a study in which, through a deep metabarcoding analysis of the 16S rRNA gene, we compared two sampling matrices, feces (F) and colonic lavage liquid (LL), in order to evaluate their accuracy to represent the complexity of the human gut microbiome. A training set of 37 volunteers was attained and paired F and LL samples were collected from each subject. A preliminary absolute quantification of total 16S rDNA, performed by droplet digital PCR (ddPCR), confirmed that sequencing and taxonomic analysis were performed on same total bacterial abundance obtained from the two sampling methods. The taxonomic analysis of paired samples revealed that, although specific taxa were predominantly or exclusively observed in LL samples, as well as other taxa were detectable only or were predominant in stool, the microbiomes of the paired samples F and LL in the same subject hold overlapping taxonomic composition. Moreover, LL samples revealed a higher biodiversity than stool at all taxonomic ranks, as demonstrated by the Shannon Index and the Inverse Simpson’s Index. We also found greater inter-individual variability than intra-individual variability in both sample matrices. Finally, functional differences were unveiled in the gut microbiome detected in the F and LL samples. A significant overrepresentation of 22 and 13 metabolic pathways, mainly occurring in Firmicutes and Proteobacteria, was observed in gut microbiota detected in feces and LL samples, respectively. This suggests that LL samples may allow for the detection of microbes adhering to the intestinal mucosal surface as members of the resident flora that are not easily detectable in stool, most likely representative of a diet-influenced transient microbiota. This first comparative study on feces and LL samples for the study of the human gut microbiome demonstrates that the use of both types of sample matrices may represent a possible choice to obtain a more complete view of the human gut microbiota in response to different biological and clinical questions.

## Introduction

The human gastrointestinal (GI) tract harbours an intricate and dynamic population of microorganisms termed gut microbiota. It includes bacteria, archaea, viruses and protists, which co-evolve and livein mutualistic cooperation with the host. About 10^14^ microbial cells colonize the human gut and outnumber by an order of magnitude the total number of human cells also providing a much higher gene complement with respect to the number of human genes (1, 2). Rapid bacterial colonization of the gut occurs during early childhood, and the intestinal microbiota gradually changes over time until adulthood, when the composition of the gut microbiota remains relatively stable (3).

Gut microbiota composition is shaped by host genetics, gender and age, diet, antibiotic usage, lifestyle, ethnicity, and living environment (4-11). Recently, various studies have revealed the importance of the gut microbiota for human health providing many benefits to the host, through a spectrum of physiological functions such as strengthening gut integrity or shaping the intestinal epithelium, harvesting energy, protecting against pathogens and regulating host immunity (12-14). The alteration in the stable composition of the gut microbiota, a condition known as dysbiosis, leads to a disruption of homeostasis with the host. Different types of dysbiosis are associated with various harmful effects on human health and long-term consequences may induce a wide range of diseases including inflammatory bowel disease (IBD), irritable bowel syndrome (IBS), diabetes mellitus, obesity, and colorectal cancer (15-20). Understanding the composition of the gut microbiota, throughout the GI tract, and the role that microbial populations can play in human intestinal health and disease can be critical for achieving a possible early diagnosis but also for the eventual development of appropriate therapeutic approaches based on the gut microbiota manipulation (i.e. prebiotics, probiotics, fecal transplantation).

In the past two decades, a multitude of studies focused on the interpretation of a “normal” and “perturbed” microbiota and almost all these studies underlined the difficulty of setting the correct sampling methods able to represent the entire and reliable composition of the gut microbiota. Hence the difficulty of making an appropriate diagnosis and consequently of choosing the most accurate therapeutic approach. Further troublings are given by the inter-individual variation of the intestinal microbiota (21,22). In fact, despite conservation at the highest taxonomic ranks, the composition of the human intestinal microbiota varies enormously from individual to individual, as regards the relative ratios of dominant phyla and the variations of genera and species linked to the individual features of the host (23-25).

Current sampling methods used to investigate the complexity of the gut microbiota are mainly based on faeces and tissue samples, obtained by endoscopy or colonoscopy such as biopsy, microdissection and luminar brush (26). Fecal samples represent the most used specimen to investigate the complexity of the human microbiome, since it represents a convenient, repeatable and non-invasive sampling method, which is able to provide sufficient biomass for gut microbiome analysis. Moreover, fecal samples provide an inexpensive method for screening a large cohort of subjects. Most of the knowledge about bacterial diversity in the human GI tract has been achieved by metagenomics analysis from fecal samples, as also reported in international projects (27,28). However, recent scientific findings showed that fecal sampling approaches may be non-exhaustive for a complete knowledge of the gut microbiota composition distributed in different sites of the GI tract (29). Just think of the different microbial composition resident in the small and large intestine both at mucosal and luminal level (30).

Biopsy, luminar brush and microdissection through endoscopy or colonoscopy are employed to better evaluate mucosa-associated microbiota. Although these procedures provide a way to investigate the mucosal microbiota composition at distinct anatomical areas of the GI tract (31-33), they show multiple flaws. For example, these sampling methods are invasive and not patient-friendly and can induce drastic changes in the intestinal microbiota due to the preliminary bowel preparation and extensive use of laxatives (34,35). Furthermore, these methods suffer from unavoidable contamination and insufficient biomass yield, are expensive, time-consuming and unsuitable for healthy controls. For these reasons, endoscopy and colonoscopy are considered non-preferential sampling procedures for gut microbiota analysis.

Recently, colonic lavage emerged as an alternative approach to assess gut microbial diversity. It consists in the injection of a large amount of fluid into the colon during colonoscopy. Although colonic lavage also requires the use of laxatives prior to sampling collection, some studies suggested that it could be a useful sampling method as it would ensure a less invasive procedure and significantly higher biomass yield than colon biopsies (36). Moreover, a new sampling approach, based on colonic lavage performed without preliminary bowel preparation and laxative, is adopted in some clinical centers and could represent another valid alternative with less impact for patients.

In this huge scenario of current sampling methods able to discriminate the gut microbial population, we performed the taxonomic characterization, through a deep metabarcoding analysis of the 16S rRNA gene (V4 region), of the gut microbiome associated with two sampling matrices, feces (F) and colonic lavage liquid (LL), in order to infer: i) their accuracy to represent the complexity of human gut; ii) intra and inter-individual variability in matched samples; iii) microbial taxa and metabolic pathways associated to both sampling matrices.

## Results

### Taxonomic characterization of gut microbiota in feces and colonic lavage liquids samples

To determine the extent and depth to which colonic LL and F sampling approaches can contribute to a broader understanding of the complexity of the gut microbiome, 37 patient-paired F and LL samples were analysed by a deep metabarcoding sequencing of the bacterial 16S rRNA gene (V4 region). About 11.7 million of paired-end reads (mean 158,596, std. dev. ± 55,230) were generated across all samples, and following the trimming, merging and denoising procedures we retained about 96.8% of initial sequences (Supplementary Figure S1). Overall, we obtained 16,105 Amplicon Sequencing Variants (ASVs) and according to the taxonomic classification we identified and removed 61 chloroplast, mitochondrial and unclassified sequences from further analysis. Moreover, we also identified 131 ASVs as contaminants and we removed them from subsequent analysis.

To investigate the detectable microbial community composition using the two different sampling methods, we performed a taxonomic analysis of fecal and colonic lavage liquid microbiomes. In the average profiles of both F and LL samples, we observed that the dominant phyla were *Firmicutes* (median F 48.77%, IQR 35.35%, 57.31% vs median LL 52.83%, IQR 43.41%, 64.85%), *Bacteroidota* (median F 38.96%, IQR 26.87%, 46.13% vs median LL 27.27%, IQR 21.53%, 37.49%) and *Proteobacteria* (median F 3.39%, IQR 1.88%, 9.38% vs median LL 5.08%, IQR 2.57%, 17.29%; Fig. 1). Phylum *Firmicutes* was the dominant one both in F and in LL samples and it was mostly composed of *Clostridia*. The phylum *Bacteroidota* was the second most represented in all samples (all belonging to class *Bacteroides*) and the phylum *Proteobacteria*, mainly owned by class *Gammaproteobacteria*, was the last most abundant in both paired samples. At phylum level, we observed that one of the paired samples of 7 subjects showed a huge increase in the relative abundance of *Proteobacteria* (differences in relative abundance for *Proteobacteria* of 30% between F and LL samples was used as threshold value). Moreover, considering 6 paired samples the increase of *Proteobacteria* occurred in LL samples (mean LL 58.62% ± 16.92 vs mean F 3.14 ± 3.76) while only for a set of paired samples in the F sample (V4-BioS-349 F 61.3% vs LL 5.18%). Such increase of *Proteobacteria* was associated with a decrease in *Bacteroidota* (mean F 52.13%, std. dev ± 20.34 vs mean LL 19.32%, std. dev ± 7.18) and *Firmicutes* (mean F 39.51, std. dev ± 12.07 vs mean LL 17.06%, std. dev ± 8.94). However, most of the F and LL paired samples showed similar communities at the phylum level.

**Figura 1.**
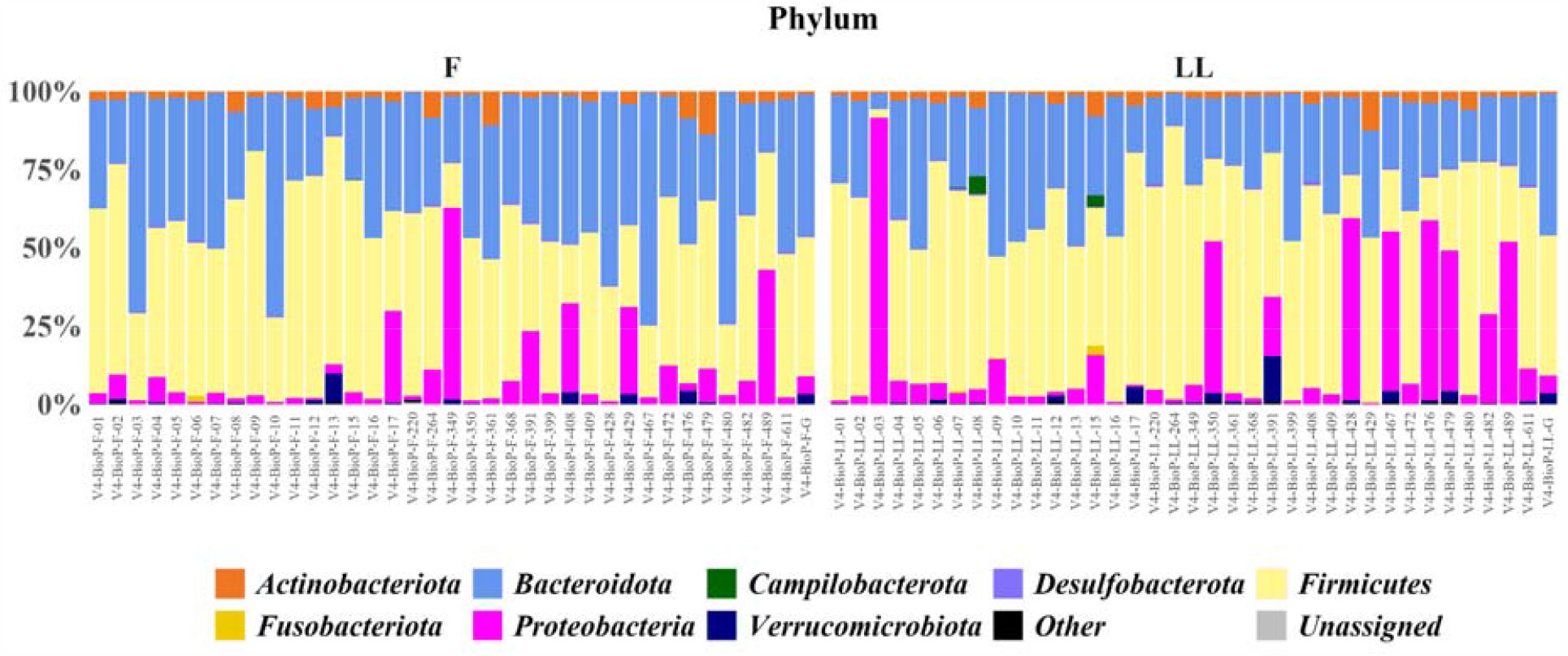
Bacteria relative abundances at the phylum level. Only taxa with a relative abundance equal or higher to 1% were shown. Rare taxa were collapsed in Other. Samples are grouped according to the sampling matrices, i.e. feces (F) and liquid lavage (LL), and ordered according to patients.

Focusing on quantitative differences at the ASVs, Family, Genus and Species level, we looked into the proportion of unique and shared taxa between F and LL sample matrices. To recapitulate the overall number of taxa observed in paired samples we used Venn diagrams. First, we set a threshold value to establish the number of reads needed to consider a taxon effectively observed and at what number of reads accuracy of observations was lost. For this purpose, we inferred the Venn proportion by using increasing threshold values (i.e. 1, 5, 10, 15, 20, 25 and 1% of the observed taxa). We verified that the proportion of common and uncommon taxa was not related to the chosen threshold, with the exception of 1% which deeply reduced the number of taxa available for comparison (Supplementary Figure S2). For the subsequent Venn data analysis, we set a threshold value equal to 5 reads. Regarding the observed ASVs, we identified 17.8% of common taxa and unique taxa, i.e. observed exclusively either in F or in LL samples (38.8% and 43.4%, respectively; Fig. 5A). At the Families, Genera and Species levels, an increase of common taxa was observed (61.7%, 53.5%, 31.6%, respectively) but again we identified unique taxa in the both specimens (Families LL 24% vs F 14.2%, Genera LL 26.7% vs F 19.8%, Species LL 38.2% vs F 30.2%; (Supplementary Figure S3)).

For a broader understanding of the accuracy of F and LL samples to represent the complexity of the human gut microbiome, we next studied paired samples for each enrolled subject by considering gut microbiome composition at each taxonomic level. We found in about 65% of the subjects the number of unique taxa in LL samples was higher than in F samples. Moreover, we found large individual differences in unique and shared taxa among all patients enrolled in our study. Indeed, the number of shared and unique taxa were extremely variable for each subject across all considered ranks. For example, looking at the Genus level, we found the number of common taxa ranged between 27 and 93 as well as the number of uncommon taxa, a range between 4 and 87 and between 3 and 62 for LL and F unique taxa, respectively (Fig. 2).

**Figure 2.**
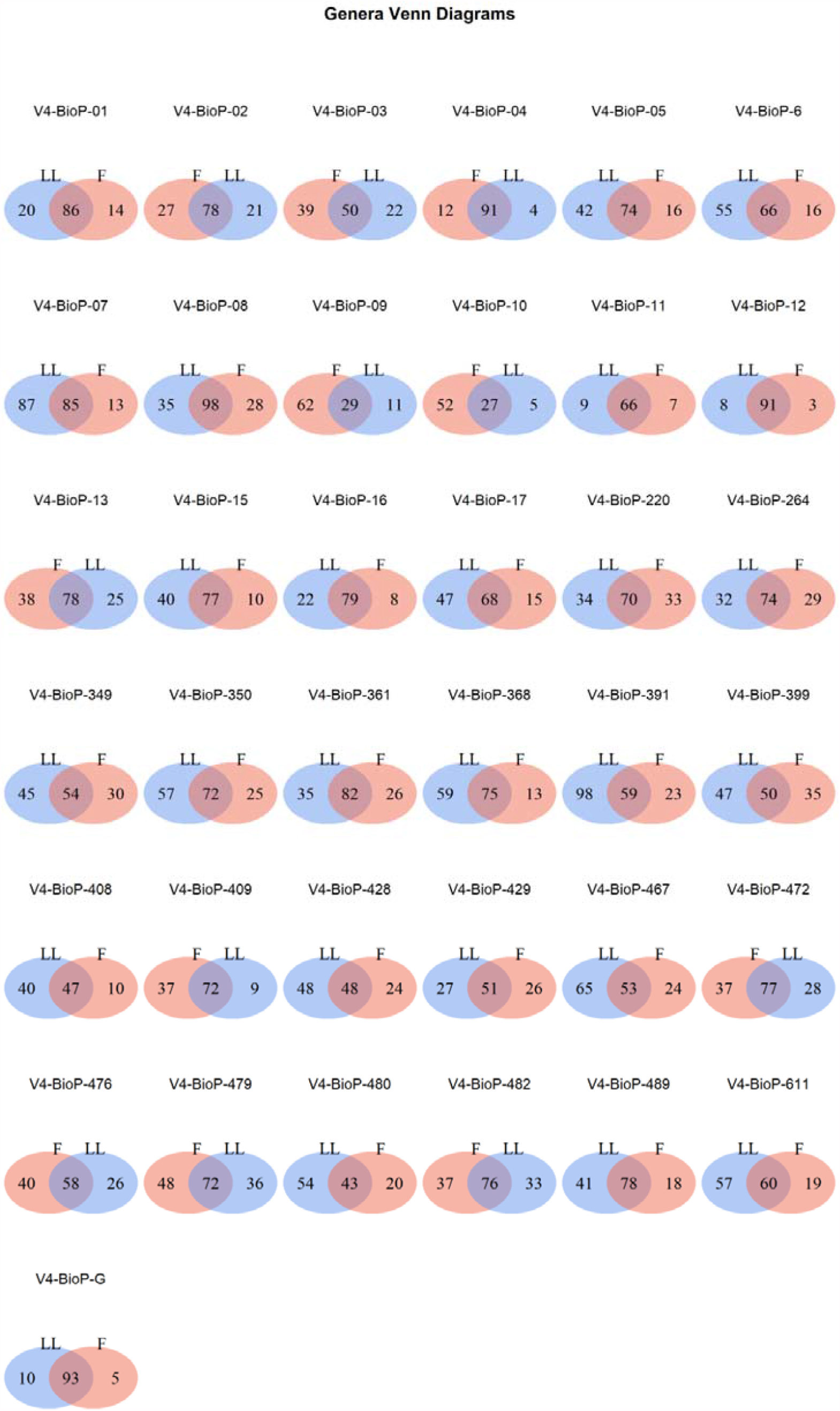
Venn Diagrams showing the percentage of shared or unique Genera observed in feces (F) and colonic lavage liquids (LL) matched samples of each patient.

### Total microbial abundance in feces and colonic lavage liquids samples

In order to evaluate gut microbial sampling differences between feces and colonic LL liquids we assessed the total microbial abundance through accurate absolute quantification of total 16S rDNA by droplet digital PCR (ddPCR). We analyzed DNA extracted from 36 paired F and LL samples. Although the DNA extraction yield was not comparable between F and LL samples because the latter are typically less concentrated than fecal DNA, we found that the total 16S copy number per ng of DNA was comparable between the two sampling approaches. As shown in Figure 3, ddPCR analysis highlighted no statistically significant differences in total 16S abundance between the two metagenomic DNA sampling, feces (mean = 846,826; st. dev = 257,116; median = 782,688; min = 463,962; max = 1,482,853; IQR = 279,960) and colonic LL fluids (mean = 963,293; st. dev. = 444,625; median = 826,583; min = 219,717; max = 2,290,411; IQR = 414,748). Some samples are an exception because they showed an LL total 16S copy number higher with respect to fecal one.

**Figure 3.**
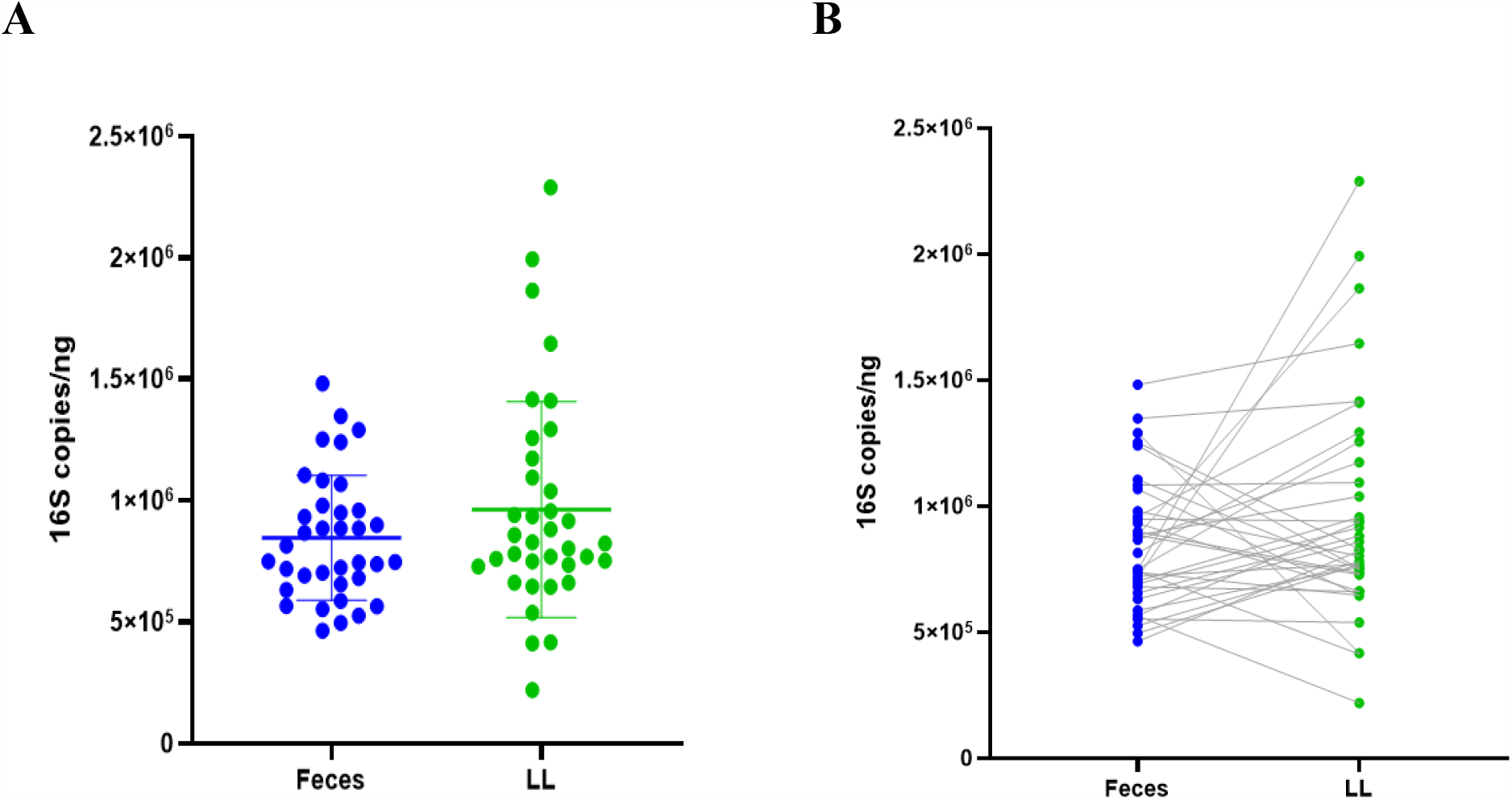
16S absolute quantification in DNA extracted from stool and LL sample matrices. Data are reported as the average of triplicate ddPCR experiments and expressed as 16S copies/ng of DNA (**A** dot-plot; **B** dumbbell plot). There are no statistically significant differences in absolute 16S copy number between feces and LL sample matrices (two-tailed t-test p-value = 0.15).

As regards the total microbial load, we can conclude that F and colonic LL sampling matrices did not present quantitative differences in the total 16S copy number and therefore were comparable by the quantitative point of view.

### Gut microbiota diversity analysis

To infer the microbial biodiversity in F and LL samples, two alpha diversity different indices, Shannon Index and Inverse Simpson Index, were calculated on the produced ASVs. No statistical difference was observed between F and LL samples by using Shannon index (paired t-test p-value 0.12, Fig. 4A) whereas the Inverse Simpson Index showed a significantly increased microbial biodiversity in the LL samples (paired t-test p-value 0.038, Fig. 4B). Following the data stratification according to gender and age (Table 5), no significant statistical changes were found between F and LL sample groups using both Shannon Index and Inverse Simpson Index (Supplementary Figure S4).

**Figure 4.**
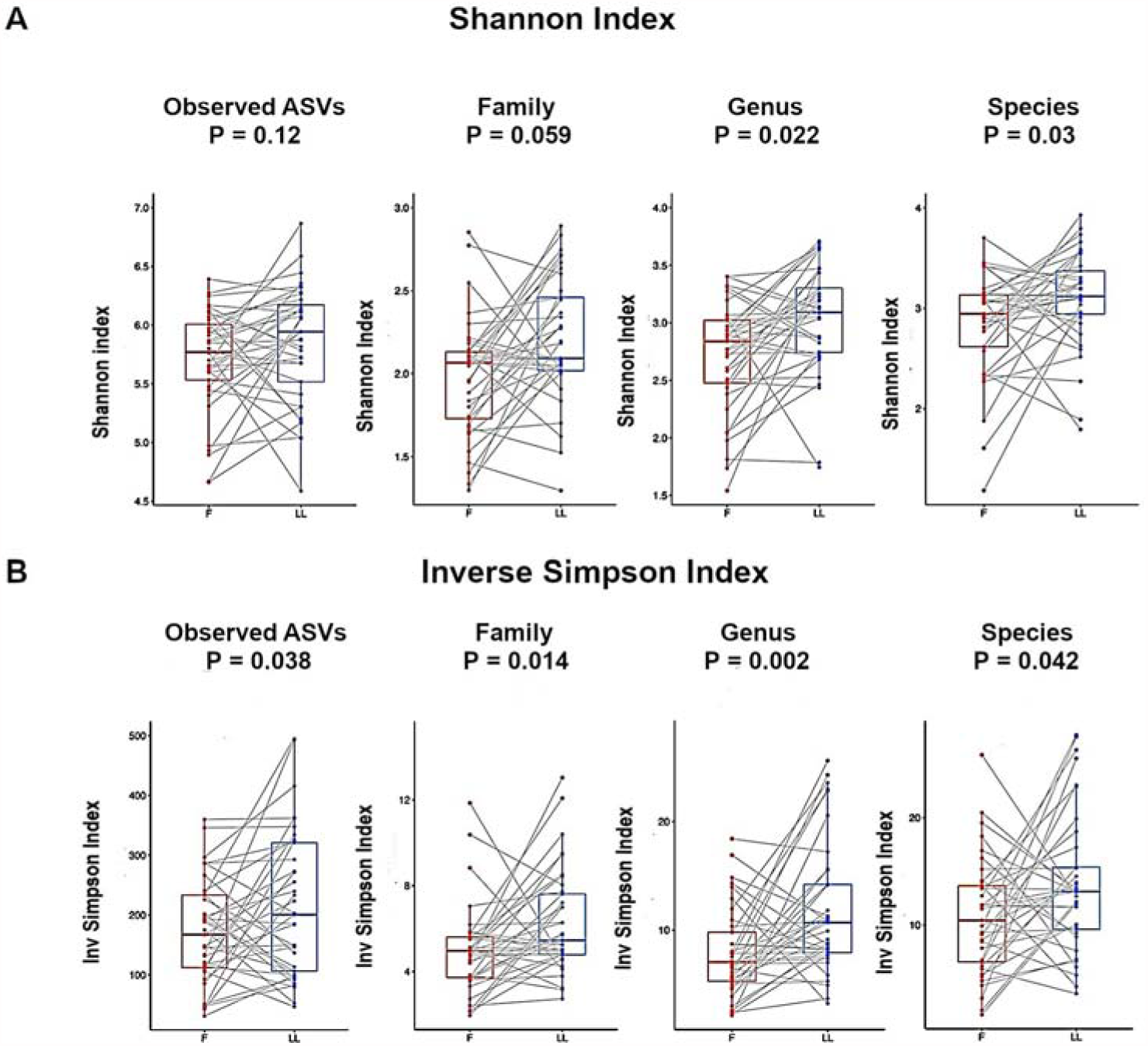
Alpha diversity comparison between faeces (F) and colonic lavage liquids (LL). The biodiversity of microbiome was measured using **A)** Shannon diversity index and **B)** Inverse Simpson’s evenness on Observed ASVs and at the Family, Genus and Species levels. Alpha diversity scores were calculated by using normalized data by rarefaction to 68,000, 67,000, 66,000 and 19,000 sequences, respectively. Each point represents the diversity score for a patient sample. Between-group variations were measured using the Student’s test (S) and the Wilcoxon tests (W) and P values for each alpha diversity measured was reported for each group. P<0.05 was considered as statistically significant.

**Table 1.** Summary results of the PERMANOVA model based on the weighted and unweighted UniFrac dissimilarity matrix.

|  | Weighted Unifrac |  | Unweighted Unifrac |  |
| --- | --- | --- | --- | --- |
|  | R<sup>2</sup> | p-value | R<sup>2</sup> | p-value |
| **Matrix** | 0.00845 | 0.199 | 0.01464 | 0.056 |
| **Gender** | 0.02063 | 0.039 * | 0.01492 | 0.041 * |
| **Age** | 0.02717 | 0.248 | 0.02833 | 0.041 * |
| **Subjects** | 0.49609 | 0.006 ** | 0.48651 | 0.001 *** |
| **Residuals** | 0.44129 |  | 0.45560 |  |

**Table 2.** Summary results of the PERMANOVA model based on the Bray-Curtis dissimilarity matrix.

|  | Family |  | Genus |  | Species |  |
| --- | --- | --- | --- | --- | --- | --- |
|  | R<sup>2</sup> | p-value | R<sup>2</sup> | p-value | R<sup>2</sup> | p-value |
| **Matrix** | 0.02844 | 0.040 * | 0.02695 | 0.016 * | 0.01689 | 0.097 |
| **Gender** | 0.02358 | 0.070 . | 0.01604 | 0.209 | 0.02111 | 0.021 * |
| **Age Group** | 0.02457 | 0.457 | 0.02527 | 0.460 | 0.03179 | 0.107 |
| **Subjects** | 0.47290 | 0.156 | 0.47378 | 0.099 | 0.49945 | 0.002 ** |
| **Residuals** | 0.45052 |  | 0.45797 |  | 0.43076 |  |

**Table 3.**
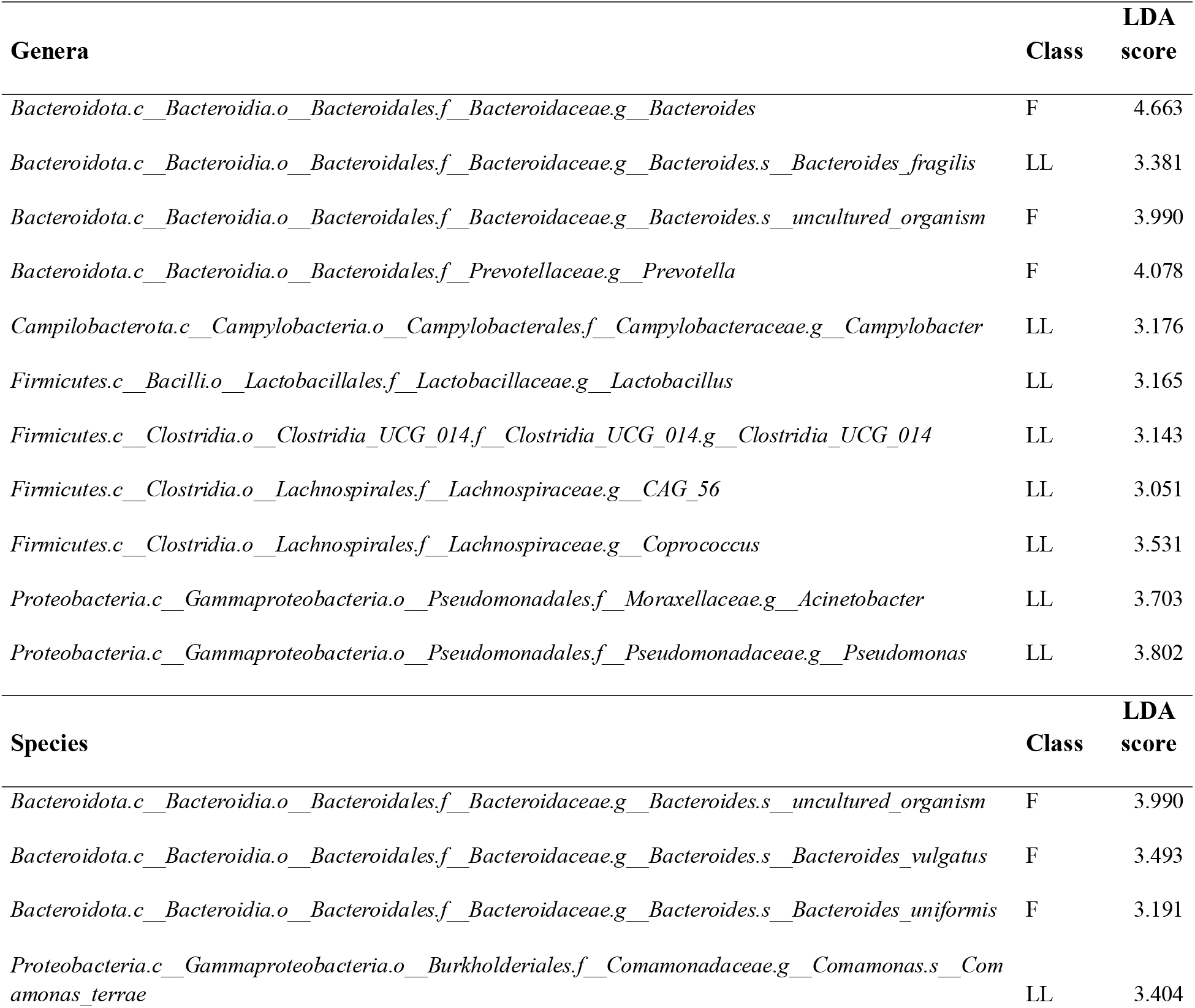
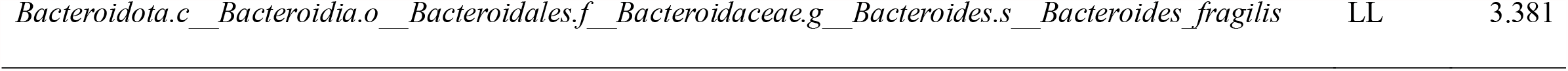
LEfSe results for Genera and Species with an LDA score ≥ 3.0.

| Genera | Class | LDA score |
| --- | --- | --- |
| *Bacteroidota.c__Bacteroidia.o__Bacteroidales.f__Bacteroidaceae.g__Bacteroides* | F | 4.663 |
| *Bacteroidota.c__Bacteroidia.o__Bacteroidales.f__Bacteroidaceae.g__Bacteroides.s__Bacteroides_fragilis* | LL | 3.381 |
| *Bacteroidota.c__Bacteroidia.o__Bacteroidales.f__Bacteroidaceae.g__Bacteroides.s__uncultured_organism* | F | 3.990 |
| *Bacteroidota.c__Bacteroidia.o__Bacteroidales.f__Prevotellaceae.g__Prevotella* | F | 4.078 |
| *Campilobacterota.c__Campylobacteria.o__Campylobacteriales.f__Campylobacteraceae.g__Campylobacter* | LL | 3.176 |
| *Firmicutes.c__Bacilli.o__Lactobacillales.f__Lactobacillaceae.g__Lactobacillus* | LL | 3.165 |
| *Firmicutes.c__Clostridia.o__Clostridia_UCG_014.f__Clostridia_UCG_014.g__Clostridia_UCG_014* | LL | 3.143 |
| *Firmicutes.c__Clostridia.o__Lachnospirales.f__Lachnospiraceae.g__CAG_56* | LL | 3.051 |
| *Firmicutes.c__Clostridia.o__Lachnospirales.f__Lachnospiraceae.g__Coprococcus* | LL | 3.531 |
| *Proteobacteria.c__Gammaproteobacteria.o__Pseudomonadales.f__Moraxellaceae.g__Acinetobacter* | LL | 3.703 |
| *Proteobacteria.c__Gammaproteobacteria.o__Pseudomonadales.f__Pseudomonadaceae.g__Pseudomonas* | LL | 3.802 |
| Species | Class | LDA score |
| *Bacteroidota.c__Bacteroidia.o__Bacteroidales.f__Bacteroidaceae.g__Bacteroides.s__uncultured_organism* | F | 3.990 |
| *Bacteroidota.c__Bacteroidia.o__Bacteroidales.f__Bacteroidaceae.g__Bacteroides.s__Bacteroides_vulgatus* | F | 3.493 |
| *Bacteroidota.c__Bacteroidia.o__Bacteroidales.f__Bacteroidaceae.g__Bacteroides.s__Bacteroides_uniformis* | F | 3.191 |
| *Proteobacteria.c__Gammaproteobacteria.o__Burkholderiales.f__Comamonadaceae.g__Comamonas.s__Comamonas_terrae* | LL | 3.404 |

**Table 4.** List of Prevalence scores describing taxa observed in F and LL samples. F: the number of time the taxa was exclusively observed in F in at least one paired sample; LL: the number of time the taxa was exclusively observed in LL in at least one paired sample; F prev: the prevalence of a taxa in F samples; LL prev: the prevalence of a taxa in LL samples; F prop: the proportion between the number of time the taxa was exclusively seen in F samples and the F prevalence; LL prop: the proportion between the number of time the taxa was exclusively seen in LL samples and the LL prevalence; Score: the difference between F and LL prevalence.

| Taxa | F | LL | F prev | LL prev | rank | F prop | LL prop | Score |
| --- | --- | --- | --- | --- | --- | --- | --- | --- |
| *d__Bacteria;p__Actinobacteriota;c__Coriobacteriia;o__Coriobacteriales;f__Coriobacteriales_Incertae_Sedis* | 5 | 14 | 19 | 28 | Family | 0,26 | 0,50 | -9 |
| *d__Bacteria;p__Firmicutes;c__Clostridia;o__Oscillospirales;f__[Eubacterium]_coprostanoligenes_group;*<br>*g__[Eubacterium]_coprostanoligenes_group;s__gut_metagenome* | 5 | 14 | 13 | 22 | Species | 0,38 | 0,64 | -9 |
| *d__Bacteria;p__Bacteroidota;c__Bacteroidia;o__Bacteroidales;f__Barnesiellaceae;g__Copro bacter* | 4 | 12 | 15 | 23 | Genus | 0,27 | 0,52 | -8 |
| *d__Bacteria;p__Firmicutes;c__Negativicutes;o__Acidaminococcales;f__Acidaminococcaceae;*<br>*g__Phascolarctobacterium* | 7 | 15 | 19 | 27 | Genus | 0,37 | 0,56 | -8 |
| *d__Bacteria;p__Firmicutes;c__Clostridia;o__Lachnospirales;f__Lachnospiraceae;*<br>*g__[Eubacterium]_ruminantium_group* | 6 | 14 | 8 | 16 | Genus | 0,75 | 0,88 | -8 |
| *d__Bacteria;p__Firmicutes;c__Clostridia;o__Lachnospirales;f__Lachnospiraceae;g__Blautia;s__metagenome* | 3 | 11 | 12 | 20 | Species | 0,25 | 0,55 | -8 |
| *d__Bacteria;p__Proteobacteria;c__Gammaproteobacteria;o__Burkholderiales;f__Burkholderiaceae;*<br>*g__Burkholderia-Caballeronia-Paraburkholderia* | 17 | 0 | 17 | 0 | Genus | 1,00 | 0,00 | 17 |
| *d__Bacteria;p__Proteobacteria;c__Gammaproteobacteria;o__Burkholderiales;f__Burkholderiaceae* | 18 | 2 | 20 | 4 | Family | 0,90 | 0,50 | 16 |
| *d__Bacteria;p__Proteobacteria;c__Gammaproteobacteria;o__Burkholderiales;f__Burkholderiaceae;*<br>*g__Burkholderia-Caballeronia-Paraburkholderia;s__Burkholderia_pseudomallei* | 14 | 0 | 14 | 0 | Species | 1,00 | 0,00 | 14 |
| *d__Bacteria;p__Firmicutes;c__Clostridia;o__Lachnospirales;f__Lachnospiraceae;*<br>*g__[Eubacterium]_fissicatena_group* | 15 | 4 | 15 | 4 | Genus | 1,00 | 1,00 | 11 |
| *d__Bacteria;p__Actinobacteriota;c__Actinobacteria;o__Propionibacteriales;f__Propionibacteriaceae;*<br>*g__Cutibacterium* | 12 | 1 | 12 | 1 | Genus | 1,00 | 1,00 | 11 |
| *d__Bacteria;p__Bacteroidota;c__Bacteroidia;o__Bacteroidales;f__Rikenellaceae;g__Alistipes;s__unidentified* | 17 | 6 | 25 | 14 | Species | 0,68 | 0,43 | 11 |
| *d__Bacteria;p__Firmicutes;c__Clostridia;o__Oscillospirales;f__Oscillospiraceae;g__UCG-002;s__gut_metagenome* | 17 | 6 | 23 | 12 | Species | 0,74 | 0,50 | 11 |
| *d__Bacteria;p__Actinobacteriota;c__Actinobacteria;o__Propionibacteriales;f__Propionibacteriaceae* | 12 | 2 | 12 | 2 | Family | 1,00 | 1,00 | 10 |
| *d__Bacteria;p__Actinobacteriota;c__Actinobacteria;o__Propionibacteriales* | 12 | 2 | 12 | 2 | Order | 1,00 | 1,00 | 10 |

**Table 5.**
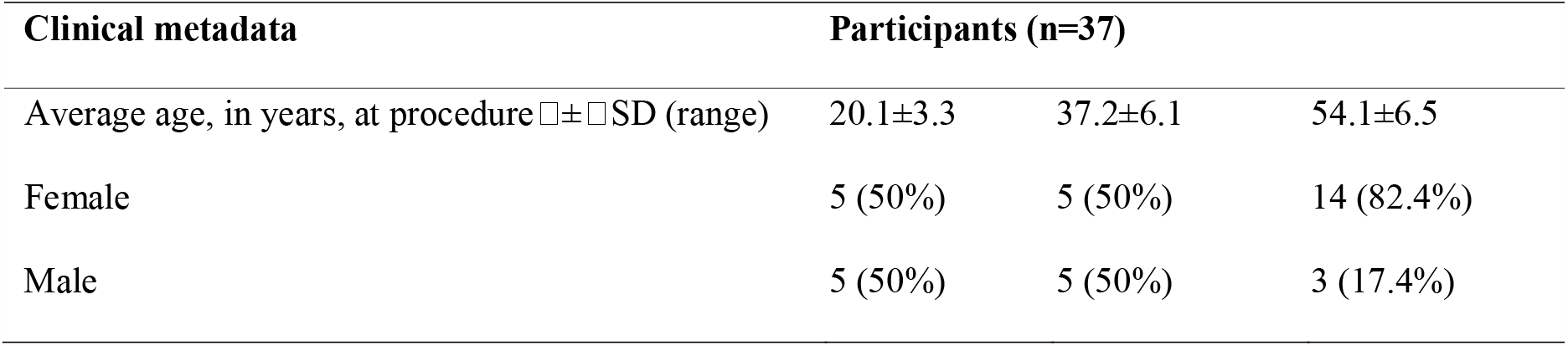
Clinical characteristics of enrolled participants.

| Clinical metadata | Participants (n=37) |  |  |
| --- | --- | --- | --- |
| Average age, in years, at procedure $\bar{x} \pm \text{SD}$ (range) | 20.1±3.3 | 37.2±6.1 | 54.1±6.5 |
| Female | 5 (50%) | 5 (50%) | 14 (82.4%) |
| Male | 5 (50%) | 5 (50%) | 3 (17.4%) |

Alpha-diversity measurements were performed at different taxonomic levels. Overall, we observed 160, 417 and 669 bacterial Families, Genera and Species, respectively, and following the rarefaction, we retained 147 Families, 397 Genera and 630 Species. Among them, we detected significant statistical differences considering the two sample matrices evaluated in this study. In particular, LL samples exhibited a greater biodiversity than F samples at all the indicated taxonomic levels, as demonstrated by the Shannon Index and the Inverse Simpson’s Index (Fig. 4A,B).

To evaluate the variability in microbial community composition among samples, we next measured beta diversity by using the weighted and unweighted UniFrac dissimilarity metrics. We used Principal Coordinates Analysis (PCoA) to visualize the differences between the samples according to the matrix of beta diversity distance. By using the weighted-UniFrac (Fig. 5A), we observed the first component (x-axis) explaining about 18.3% of the total variability and the second one 7.4% (y-axis). In the unweighted-UniFrac PCoA (Fig. 5B), we detected the first component (x-axis) explaining about 4.2% of the total variability and the second one 4.1% (y-axis). In both PCoA plots of the F and LL samples dataset we observed that the paired samples were partially clustered according to the individual-subject underlining the strong individual uniqueness of the gut microbiome.

**Figure 5.**
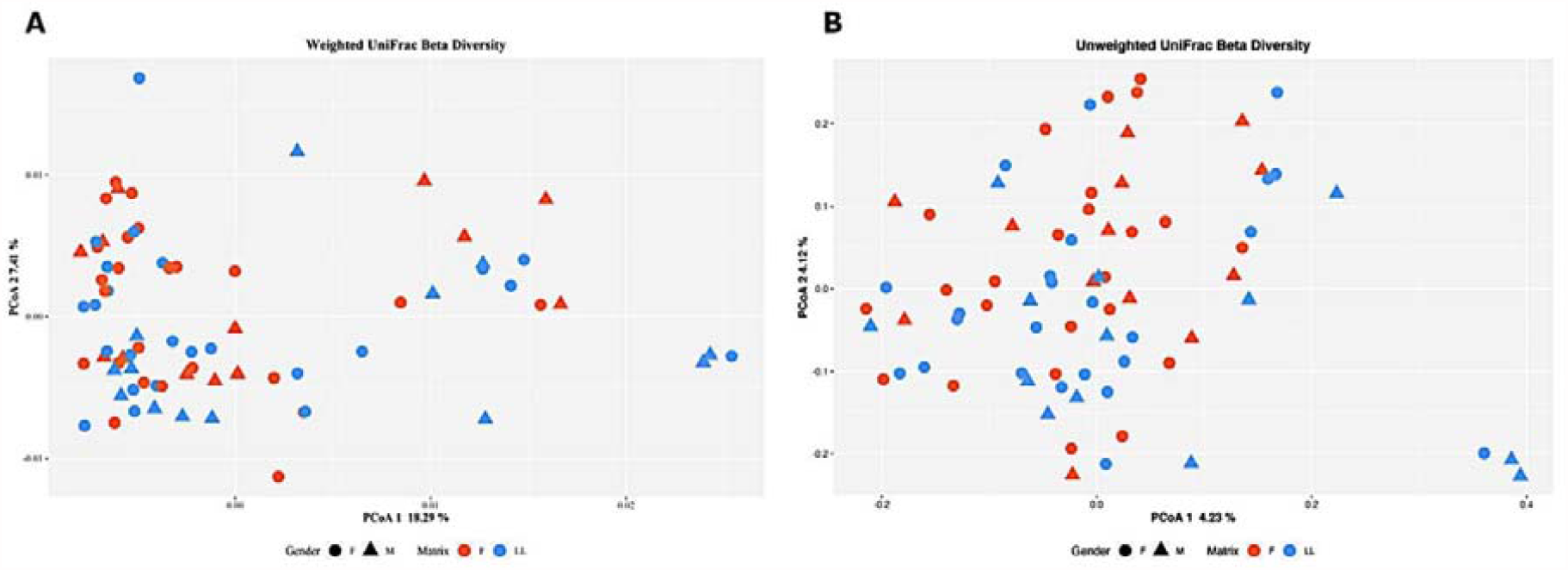
PCoA plot based on the weighted and unweighted UniFrac dissimilarity matrix. We performed the PERMANOVA analysis (Table 1) based on the measured dissimilarity matrices taking into account the patient-paired F and LL samples, gender, age group and subjects. According to the PERMANOVA analysis results for the weighted UniFrac metric, the inter-individual variability of Subjects and Gender groups significantly explained about 49.6% and 2% of the observed variability, respectively. None of the other variables obtained a significant result. The PERMANOVA applied to unweighted UniFrac metric, confirmed that inter-individual diversity explained the largest part (48%) of the observed variability and that Gender and Age groups significantly explained a smaller part of the observed data (1.5% and 2.8%, respectively; Table 2).

The Families, Genera and Species rarefied counts were used to measure beta diversity by using the Bray-Curtis metrics. Once again, the PCoA showed a clear clustering based on subjects (Supplementary Figure S5) but no clustering based on sampling approaches.

Indeed, in the PERMANOVA model, built by using sampling matrices, gender, age, and subjects as covariates, subjects explained about the 50% of the observed variability at families, genera and species level.

### Microbial taxa and metabolic pathways correlation

To identify microbial taxa and metabolic pathways associated to the sampling matrices under evaluation, we performed a Linear discriminant analysis Effect Size (LEfSe) analysis. All in all, 169 taxa and 35 metabolic pathways were associated with the sample types. In particular, we identified 128 taxa associated with the LL sample groups that were not observed or were in much lower amounts in the fecal samples. A further 41 taxa were correlated to the F samples since they were exclusively observed or present in higher amounts than the LL samples. Taking into account the taxa assigned at the Genus level, we identified *Prevotella, Bacteroides* and *Sellimonas* genera associated with F samples and 26 genera associated to LL samples (Supplementary Figures S6). Among them, *Lactobacillus, Campylobacter, Coprococcus, Acinetobacter, Pseudomonas* and two genera belonging to *Clostridia* class were associated to LL samples with LDA score ≥ 3 (Table 3). At species level, we observed 68 taxa associated with LL samples, mostly belonging to *Clostridia, Bacteroidia* and *Gammaproteobacteria* classes. On the other hand, 31 species were associated to F samples, above all belonging to *Clostridia* and *Bacteroidia* classes. Interestingly, considering only the taxa with LDA score ≥ 3, we observed that *Bacteroides vulgatus* and *Bacteroides uniformis* were associated to F samples, while *Bacteroidetes fragilis* was strongly related to LL samples (Table 3). A heat map of the retained 169 taxa revealed that F and LL samples did not form discrete clusters by sample types, age and gender groups indicating an inter-individual samples separation versus an intra-individual paired samples separation (Supplementary Figures S7).

To assess the functional differences in the gut microbiome detected in the F and LL samples, we predicted the functional content based on the 16S rRNA data by using PICRUSt, which revealed a significant overrepresentation of 22 and 13 pathways in F and LL samples, respectively. Interestingly, the 13 metabolic pathways associated with LL samples mainly occurred in bacteria belonging to the phylum *Proteobacteria*. Figure 6 shows the heat map of the 35 pathways associated to each type of sample. he 7 subjects previously described with increased *Proteobacteria* levels in one of the paired samples (6 LL samples and 1 F sample) clustered together and showed a different profile from the other samples.

**Figure 6.**
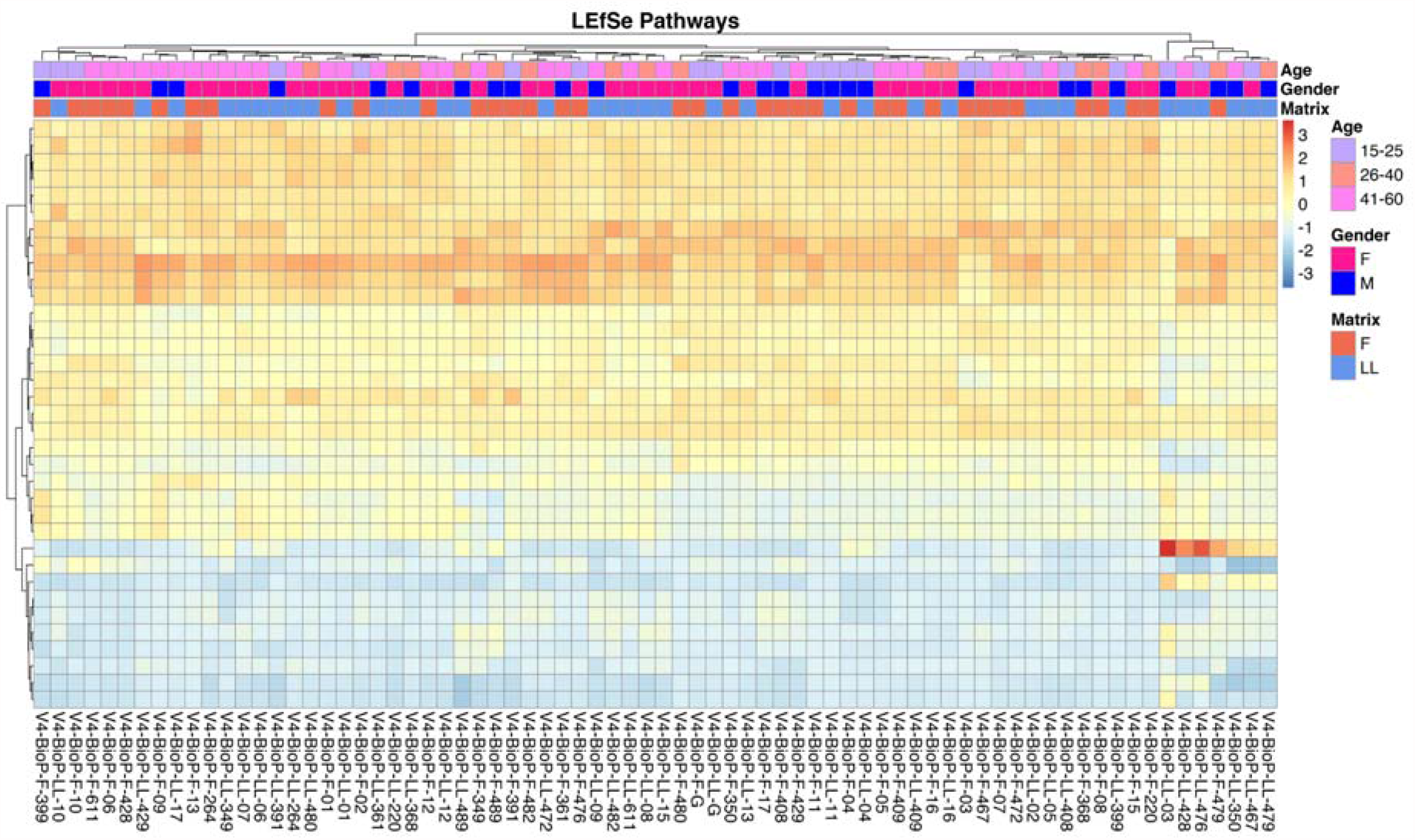
Heat map representing the LEfSe selected pathways.

To evaluate the taxa present in F but not in LL matched samples and vice versa, scatter plots were generated for each taxonomic level (Fig. 7A,B). To verify whether the taxa observed exclusively at least in a paired sample were shared among different subjects, we used the prevalence score (Score = F prevalence - LL prevalence) which allowed us to quantify how much a taxon is most observed in F or LL samples. The higher the number of samples in which a taxon is observed, the closer to zero the score. In both scatter plots for F and L samples we noticed that although most taxa were shared between the sample matrices (score close to zero), there were taxa that prevailed in the F samples and other taxa in the LL samples.

For the F samples, scores higher than 15 were observed for the family *Burkholderiaceae* (observed in 20 F and 4 LL samples, 18 times only in F and 2 times only in LL samples, score 16) and genera group *Burkholderia-Caballeronia-Paraburkholderia* (observed 17 times only in F samples, score 17; Table 4). The highest scores obtained for the LL samples was -9 for the family *Coriobacteriales_Incertae_Sedis* (observed in 19 F and 28 LL samples, 5 times only in F and 14 only in LL, score -9) and -9 for the unidentified species *[Eubacterium]_coprostanoligenes_group;s__gut_metagenome* (observed in 13 F and 22 LL samples, 5 times only in F and 14 only in LL, score -9; Table 4).

To investigate the impact of the taxa observed exclusively in one type of sample in each subject, we measured the difference between the ASVs relative abundances in F and LL samples by considering only ASVs supported by at least 5 reads. We showed the distribution of the difference in ASVs relative abundances among F and LL samples as boxplot (Fig. 7C). An average of about 69.58% (median 70.62%, min 43.28%, max 78.07%, IQR 4.61%) of ASVs was enclosed in the whisker boxplot (i.e. IQR - 1.95*IQR IQR + 1.95*IQR). This value represented the ASVs whose relative abundance was slightly different or equal between the two sample types. Although these ASVs dominated the observed distribution, they accounted for 53.14% and 22.32% of taxa abundances on average in F and LL samples, respectively (F median 54.10%, min 19.52%, max 74.78%, IQR 19.63%; LL median 19.68%, min 6.37%, max 51.20%, IQR 10.1%).

**Figure 7.**
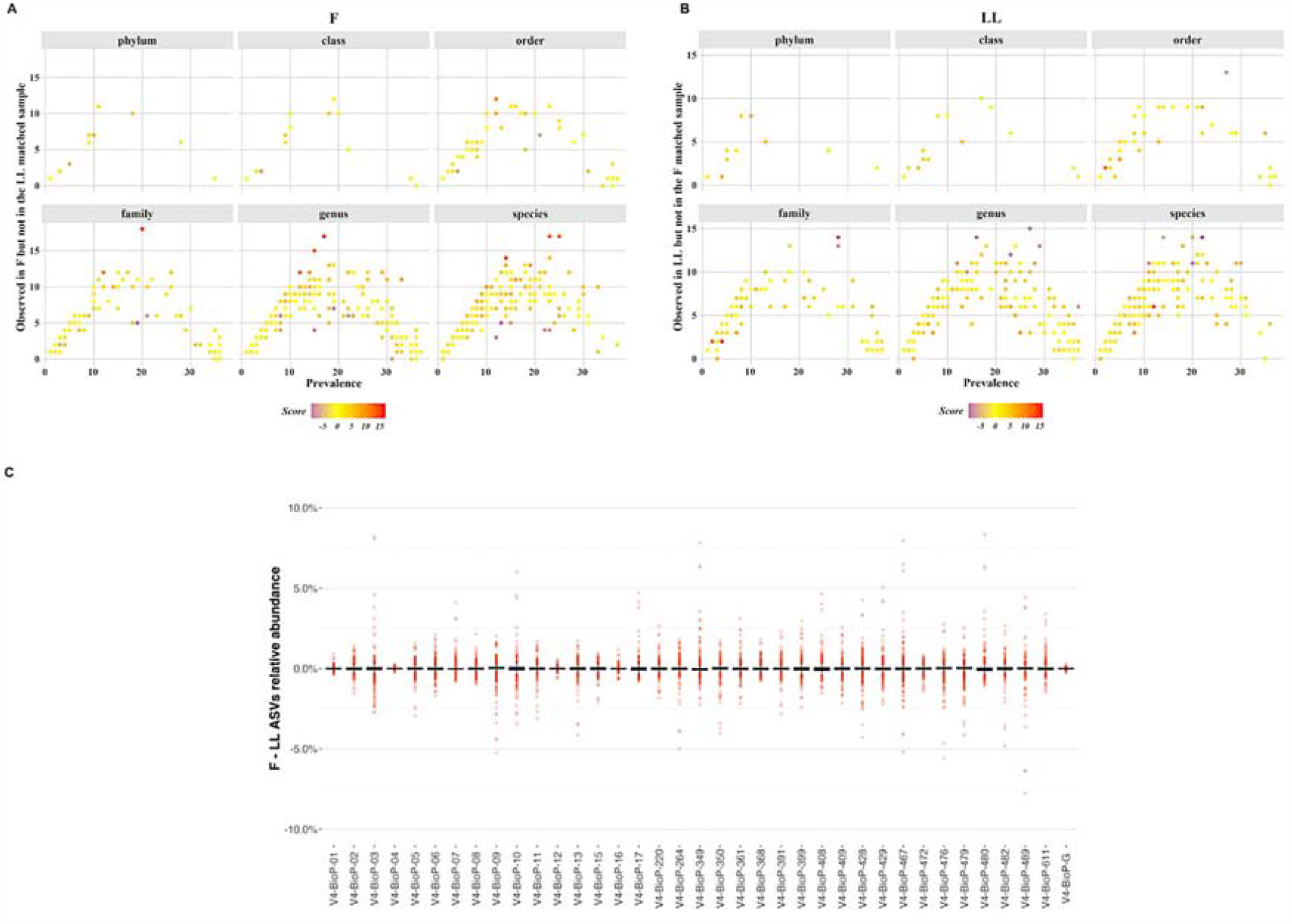
**A**. Scatter Plot representing the taxa prevalence in F samples (x-axis) against the number of times the taxa is exclusively observed in F (y-axis). Point color is equal to the difference among F and LL prevalence (score). **B**. Scatter Plot representing the taxa prevalence in LL samples (x-axis) against the number of times the taxa is exclusively observed in LL (y-axis). Point color is equal to the difference among F and LL prevalence (score). **C**. Distribution of the difference in ASVs relative abundances among F and LL samples.

Furthermore, considering the outlier values, we observed that both the number of outliers and their divergence between the two sampling procedures were strictly related to the subject.

These data confirmed what we previously observed by comparing taxa prevalence with the number of times they were exclusively observed in F or LL samples (Fig. 7A, B). Moreover, the subjects in which larger differences were observed between the paired F and LL samples corresponded to the 7 subjects where we found a higher variation in Phyla representation (Fig. 1A).

## Discussion

Understanding the role of the gut microbiota in the well-being of the human host is a constantly moving field. Although several studies have shown that the gut microbiota offers many benefits to the host and that dysbiosis actively attends in the development of a wide variety of diseases, the choice of a universal sampling method for studying the gut microbiome still poses an issue.

Currently sampling procedures for gut microbiome investigation are mainly focused on the collection of feces. Significant evidence makes it clear that stools can not entirely catch all the biodiversity present along the GI tract. Biopsy of different intestinal portions revealed specific microbial niches, exhibiting some differences between colon mucosal and stool profiles. In recent years, colon lavage was proposed as a new approach for gut microbiome inquiry able to provide observations similar to colon biopsy but less invasive than the latter. As a biopsy, the collection of colon lavage samples is performed in patients needing colonoscopy and consequently the study of the human microbiome extended to large cohorts with this alternative sampling method remains a challenge. Moreover, in the colon lavage procedure, bowel preparation and laxatives usage are required the day before treatment.

This study attempts to seek a suitable alternative sample type, namely (colonic) lavage liquid after colon cleansing treatment for gut microbiome study. We compared (colonic) lavage liquid samples to commonly used fecal samples with the intent of highlighting likeness and differences in the gut microbial population. The advantages of this alternative sampling method consist in avoiding bowel preparation and laxatives usage during colon cleansing treatment. However, the colon cleansing technique is invasive for the patient as it takes time and requires cleaning the entire colon from the rectum to the caecum using large amounts (18– 20 liters) of physiological, filtered and temperature-regulated warm water, which entered and exited the colon through tubes connected to a rectal catheter. Hence, the difficulty of recruiting volunteers for treatment. In our study, we analyzed a training set of 37 volunteers who were asked to first collect the fecal sample and then undergo the colon cleansing treatment.

We investigated the gut microbiome using a deep metabarcoding approach that selectively amplifies the hypervariable region V4 of the bacterial16S rRNA gene. We first computed taxonomic analysis in matched samples with the purpose to verify the accuracy of F and LL to represent the complexity of human gut and to check the intra-subject taxonomic variability due the two different sampling approaches.

Most of the inferred organisms (>95%) in both F and LL sample matrices were members of the *Bacteroidetes, Firmicutes*, and *Proteobacteria* phyla, which is concordant with other studies on the human gut microbiome (29,37). The Gram-negative bacteria belonging to the *Bacteroidetes* phylum are common, abundant, and diverse within the human GI tract. They perform the metabolic conversion essential for the host, often related to the degradation of proteins or complex carbohydrates (38). In particular, the majority of the GI *Bacteroidetes spp*. identified in our study belongs to the *Bacteroidaceae, Prevotellaceae* and *Rikenellaceae* families which produce succinic acid, acetic acid and propionic acid as the main final products (38). Interestingly, our results showed that *Prevotella* and *Bacteroides* genera are associated to F samples. Of great interest is the ratio Prevotella /Bacteroides to control an unbalanced food tendency. A recent review by Gabriela Precup and Dan-Cristian Vodnar analyzed the possibility of using *Prevotella* as a potential biomarker for homeostasis or disease state through its metabolite signature (39). Several species of *Bacteroides* are dominant bacteria able to provide nutrition and vitamins to the host and other intestinal microbial residents. Depending on the location in the host, some species of *Bacteroides* may be beneficial in the intestine but pathogenic opportunistic in other locations of the body. *Bacteroides vulgatus* and *Bacteroides fragilis* were isolated from patients suffering from Crohn’s disease, in addition *Bacteroides fragilis* was associated with intra-abdominal abscesses, appendicitis, and inflammatory bowel disease. As thoroughly reviewed by us in Marzano et al., 2021 (19) we highlighted the interaction between some species, including *Bacteroides fragilis* and the colorectal cancer, suggesting the identification of the gut microbiota biomarkers with diagnostic, prognostic or predictive significance, as well as the the possibility of intestinal microbiota modulation to prevent cancer or enhance the effect of specific therapies. In this study, we found *Bacteroides vulgatus* and *Bacteroides uniformis* were associated to F samples. On the other hand, *Bacteroidetes fragilis* was strongly related to LL samples, probably due to its interaction and connection with intestinal mucin (40). These findings stress the importance to select the correct matrix for biomarker research and the use of both matrices, fecal and (colonic) lavage liquid, could be the ideal choice for an in-depth study of the intestinal microbiome.

The second component of the GI microbiota that we found abundant in both F and LL samples was the *Firmicutes* phylum which is known to dominate the butyrate biosynthesis (mainly due to *Faecalibacterium prausnitzii*) (41). Within this phylum, the most abundant GI microorganisms were members of the *Ruminococcaceae* and *Lachnospiraceae* families, belonging to the *Clostridia* class. The prominent representation of these functional groups among the subjects analyzed was in agreement with the role of these bacteria in the maintenance and protection of normal colonic epithelium and in the production of regulatory T (Treg) cells (42,43). Of note, among *Firmicutes*, our data showed that *Lactobacillus* was particularly associated with the LL samples. Several species and strains of *Lactobacilli*, including *Lactobacillus acidophilus, Lactobacillus casei, Lactobacillus rhamnosus*, and *Lactobacillus helveticus*, were extensively studied in the prevention of human and diseases (44). Nowadays, some of these species are widely used as probiotics. It is obvious that the characterization of the gut microbiome represents a critical aspect for an accurate personalized diagnosis and therapy of the host. As regards bacteria belonging to the *Proteobacteria* phylum, they are commonly detected in the human microbiota and this group of Gram-negative bacteria is particularly various, although not very abundant in the GI tract. In fact, generally all *Proteobacteria* account for about 4% of the total gut microbiota in healthy people (45). Several studies showed that an increased prevalence of the *Proteobacteria* phylum is a marker of dysbiosis (45,46). Our results demonstrated that microorganisms belonging to this phylum are more observed in lavage fluid than in faeces, thus making the LL-sampling method probably more informative in identifying possible markers of dysbiosis.

Our preliminary diversity analysis confirmed that both sampling approaches are capable of representing a good approximation of the human gut microbiome and suggested that both F and colonic LL samples could be equally informative for the study of the human microbiome.

However, although the microbiomes of the paired samples F and LL in the same subject hold overlapping taxonomic composition, our Venn diagrams revealed that specific taxa were observed exclusively in LL samples, such as species belonging to *Clostridia, Bacteroidia* and *Gammaproteobacteria* classes, as well as other taxa belonging to *Clostridia* and *Bacteroidia* classes that were detectable only in stool.

To verify whether the observed variability in taxonomic composition among paired samples is related to a different overall microbial load, we performed an accurate absolute quantification of total 16S rDNA by droplet digital PCR (ddPCR). We confirmed that no statistically significant differences were found among the F and LL matrices.

We next inferred the diversity analysis and observed that LL samples showed a greater biodiversity than F samples. In particular, measurements of alpha diversity at ASVs, Family, Genus and Species level were higher in the LL samples.

Furthermore, when we focused on paired F and LL samples collected from the same subject, we observed that the gut microbiota composition was highly individual-specific, in fact, as confirmed by beta diversity analysis, the inter-individual variability seemed to be the major cause of the observed diversity among the samples. This could most likely be due to confounding factors, such as diet or stressful conditions, lifestyle and so on, which may affect the gut microbial composition (5,6,8,47).

Finally, we identified 128 taxa associated with LL sample groups and further 41 taxa related to F samples as observed exclusively or present in greater amounts.

We may assume, further deepening it in future studies, that LL-associated microbes are not easily detectable in the feces probably considering their propensity to adhere to the gut mucosal surface as members of the resident flora in eubiotic condition, which is generally altered in the initiation of inflammatory processes during dysbiosis (43). On the contrary, F-associated microbes, usually belonging to the transient microbiota, could be related to the anatomical conformation of the final GI tract or to the last phases of the digestive processes, therefore more influenced by diet. Consequently, it could be inferred that the use of both types of sample matrices may represent a possible choice to obtain a more complete view of the human gut microbiota. However, this hypothesis needs to be confirmed by larger and independent cohorts of patients and healthy subjects. A drawback of our study is indeed the small sample size, due to the inherent difficulty of recruiting volunteers for the LL procedure. However, since there were no prior comparative studies on F and LL samples for the study of the human gut microbiome, we decided to initially conduct a preliminary study. Since our results are very encouraging, a more structured study with a large sample size is now needed.

Nowadays, new therapeutic approaches, known as personalized medicine, have opened a new window in modern medicine, and the accurate individual-specific characterization of the gut microbiome seems to be one of the most interesting aspects for future research aimed at an appropriate treatment of diseases (48,49). In this preliminary study, we show the usefulness of an in-depth analysis of gut microbiome through faeces and colonic liquid lavage as both matrices can represent two sides of the coin for investigating the human gut microbiome and unraveling individual-specific differences.

## Methods

### Subject recruitment and sampling approaches characteristics

A total of 37 participants were recruited from PoliSmail - Integrated Biomedical Services for Allergic and Immunological Diseases (Lecce), following written informed consent. Age and gender were considered as clinical characteristics of enrolled participants (Table 5).

Two different sampling approaches were performed. The first was based on the usual collection of the fecal sample (F), the other on a new method represented by the collection of the (colon) lavage liquid (LL) after about 20-30 minutes from a cleansing of the entire colon with physiological, filtered and temperature-regulated warm water (18–20 litres). Paired samples, i.e. feces and (colon) lavage liquid were collected from all the participants recruited in the study.

### Sample collection

Two samples per participants, 1g of fresh stool sample (F) and about 1mL of colonic lavage liquid (LL), were collected and preserved in two different DNA/RNA Shield™ Fecal Collection tubes (Zymo Research, USA) and stored at 4 °C until further analysis. In particular, fresh stool samples were collected from each participant prior to the colon cleansing treatment resulting in the lavage liquid collection. The practice involved cleaning the entire colon from the rectum to the caecum using large amounts (18–20 litres) of physiological, filtered and temperature-regulated warm water, which entered and exited the colon through tubes connected to a rectal catheter. After 30 minutes, (colonic) LL was collected.

### DNA Extraction From feces and colonic lavage liquid and 16S rRNA Amplicon Sequencing

Genomic DNA was isolated from F and LL samples using the FastDNA™ Spin Kit for Soil (MP Biomedicals, Santa Ana, CA, USA) according to the manufacturer instructions. A 40 s bead-beating step at speed 6 was executed on the FastPrep Instrument (BIO 101, Carlsbad, Canada). Genomic DNA was eluted in a 100-μL volume of sterile water and stored at -20 °C. DNA quality was evaluated using the Agilent TapeStation 2200 System (Agilent, Santa Clara, CA, USA) with Genomic DNA ScreenTape assay (Agilent, Santa Clara, CA, USA). The DNA Integrity Number (DIN) was assigned to each sample by applying Agilent 2200 TapeStation software (controller version A.01.05). Then, quantitative fluorimetric analysis was carried out using Quant-iTTM PicoGreen® dsDNA Assay Kit (Invitrogen, Carlsbad, California) on a NanoDrop 3300 Fluorospectrometer (Thermo Fisher Scientific, Waltham, MA, USA).

The V4 hypervariable region of the bacterial 16S rRNA gene was chosen as the target for prokaryotic identification and the amplicon library was achieved according to established procedures (50,51). The V4 region was amplified using 0.5 ng of DNA extracted from F and LL and the universal primer pairs 515F and 806R (underlined nucleotides in the following sequences) designed to contain from 5′ to 3′ ends the transposon Nextera’s sequences (Nextera DNA sample preparation guide, Illumina): 515F, 5′-TCGTCGGCAGCGTCAGATGTGTATAAGAGACAG/GTGCCAGCMGCCGCGGTAA-3′, and 806R, 5′-GTCTCGTGGGCTCGGAGATGTGTATAAGAGACAG/GGACTACHVGGGTWTCTAAT-3′.

All PCRs were done in the presence of a non-template-control PCR reaction (negative control). The PCR products were purified using the AMPure XP Beads at a concentration of 0.8 vol/vol (Agencourt Bioscience Corporation, Beverly, Massachusetts) employing the Hamilton Microlab STAR Liquid Handling System (Hamilton Company, Reno, NV, USA). Obtained amplicon libraries (around 420 bp long) were then quantified by the Quant-iTTM PicoGreen® dsDNA Assay Kit (Invitrogen, Carlsbad, CA, USA) using the NanoDrop™ 3300 Fluorospectrometer (Thermo Fisher Scientific, Waltham, MA, USA).

Equimolar ratios of the purified amplicons were pooled and subjected to 2 × 250 bp paired-end sequencing on the MiSeq platform (Illumina, San Diego, CA, USA). To increase the genetic diversity, as required by the MiSeq platform, 25% of phage PhiX genomic DNA library was added to the mix and co-sequenced.

### ddPCR experiments

Digital PCR was performed to determine the total number of 16S copies by using universal primers targeting the V5–V6 regions of 16S rDNA (primer sequences: forward, B-V5□:□5′-ATTAGATACCCYGGTAGTCC-3′; reverse, A-V6, 5′-ACGAGCTGACGACARCCATG-3′), according to established procedures (52).

Absolute quantification was performed using QuantaSoft version 7.4.1 software (Bio-Rad, Hercules, CA, USA,) and the negative/positive thresholds were set manually, excluding samples with a number of droplets <10□000. Output results were expressed in 16S copies µl-1.

### 16S rRNA Sequencing processing

Raw 16S rRNA sequences data were quality checked by using FastQC (Available online: http://www.bioinformatics.babraham.ac.uk/projects/fastqc/) and multiQC (53). Illumina adapters and PCR primers were removed from raw reads by applying cutadapt (54) and the retained PE reads were merged by using PEAR (55). Then, the obtained merged amplicons were denoised into ASVs (Amplicon Sequence Variants) (56) by applying DADA2 (version 1.10.1) (57). ASVs were taxonomically annotated by using BioMaS (Bioinformatic analysis of Metagenomic amplicons) (58) and using the release 138 of the SILVA database (59). Contaminant ASVs were identified by using decontam (60) and those assigned to chloroplast or mitochondria were removed from subsequent analysis. The retained ASVs were multi-aligned by applying MAFFT (61) and the obtained multiple sequence alignment (MSA) was masked by using DECIPHER (62,63). A maximum-likelihood (ML) phylogenetic tree based on the masked MSA was obtained in Fasttree 2 (64). The R packages phyloseq (1.26.1) (65) and vegan (2.5-6) (66) were used to measure alpha and beta diversity. For this purpose, ASVs counts were normalized by using rarefaction (depth values settled to 68,000). In particular, the Shannon and Inverse Simpson indexes were used as measures of alpha diversity (i.e. intra-sample diversity), and the unweighted and weighted UniFrac (67) dissimilarity matrices were used to measure the beta diversity (i.e. inter-sample diversity). Statistical differences in alpha diversity indexes were measured by using the Student’s t-test (S) and the Wilcoxon test (W). The PERMANOVA (Permutational analysis of variance) was measured to infer the explained variability in beta diversity data by applying 999 permutations. Prediction of metagenome metabolic pathways was performed by using PICRUSt2 (68) by using the MetaCyc database (69) as reference. Alpha and Beta diversity inferences were also performed on data aggregated at family, genus and species level and rarefied by using 67,000, 66,000 and 19,000 sequences, respectively. The same metrics used on rarefied ASVs counts were applied except for beta-diversity analysis which was based on the Bray-Curtis dissimilarity matrices.

Finally, the association between taxa and metabolic pathways and the sampling matrix was performed by using LEfSe (70). Taxa and metabolic pathways with an LDA score ≥ 2.0 were retained.

## Author contributions statement

Must include all authors, identified by initials, for example: G.P. and M.Mi. conceived the experiment(s), E.P., B.F., M.D.R., E.N., A.O., C.M., M.Ma., S.B., D.M., Mar.Mi., I.V. and A.D. conducted the experiment(s), E.P., B.F., M.D.R., M.Ma., A.D., and G.P. analysed the results. All authors reviewed the manuscript.

