## Supplementary Figures for "Natural and after colon washing fecal samples: the two sides of the coin for investigating the human gut microbiome"

**SUPPLEMENTARY MATERIAL**

**“Taxonomic characterization of gut microbiota in feces and colonic lavage liquids samples.”**

**Figure S1**

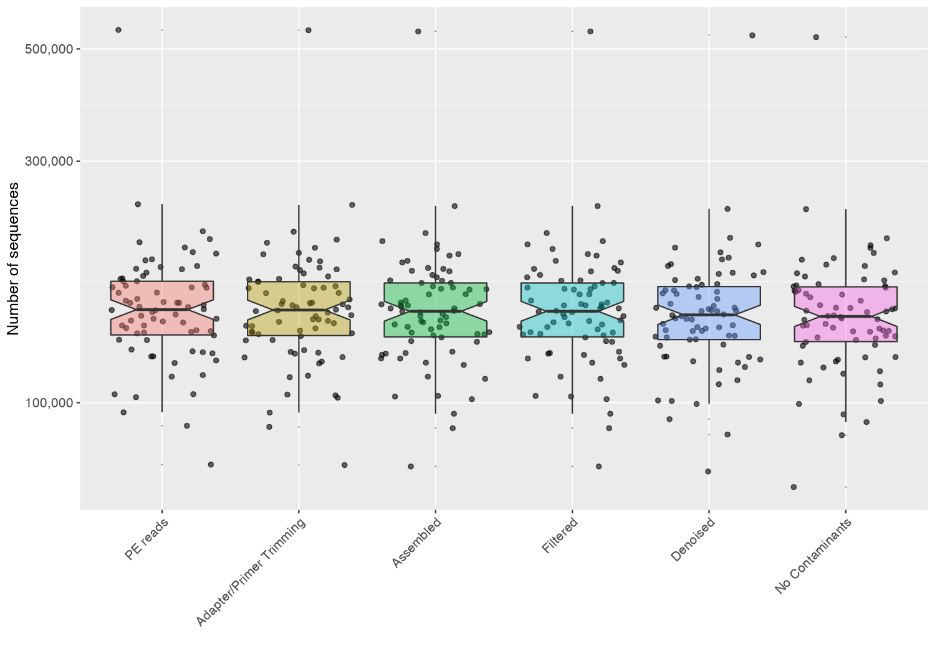

**Figure S1. Boxplot representing the per sample sequences counts from raw data to denoised one.** In particular: (i) PE reads: number of produced PE reads; (ii) Adaptor/Primer Trimming: number of PE reads retained after the Illumina adapter and PCR primers trimming; (iii) assembled: number of merged PE reads; (iv) Filtered: number of merged sequences passing the DADA2 quality filter; (v) Denoised: number of denoised sequences; (vi) No Contaminants: number of retained sequences following the removal of mitochondrial, chloroplast and unclassified ASVs.

**Figure S2**

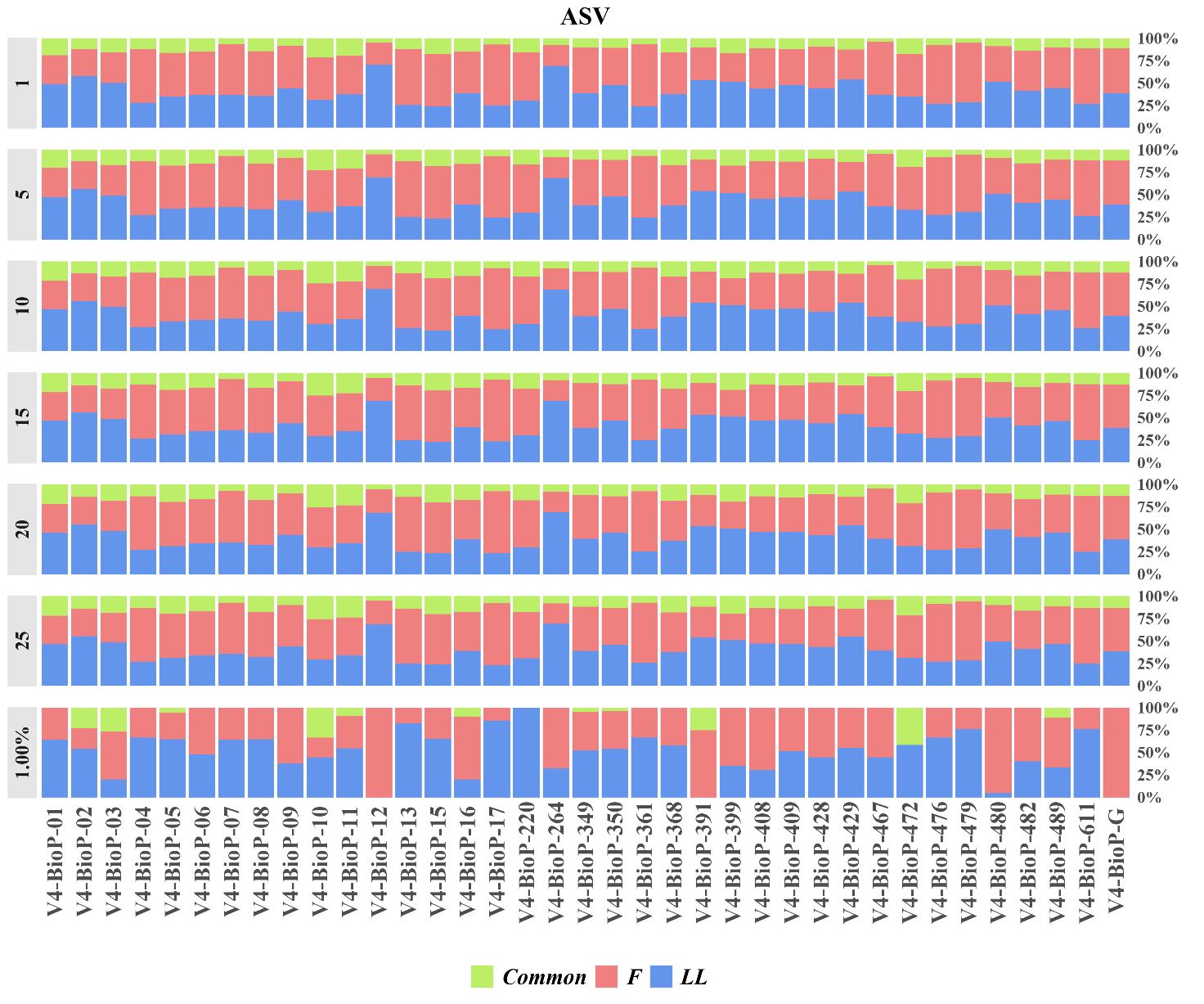

**Figure S2. Relative proportion of observed ASVs.** Percentage of Common (green) and uncommon ASVs exclusively in F samples (red) or in LL (blue) are reported.

**Figure S3**

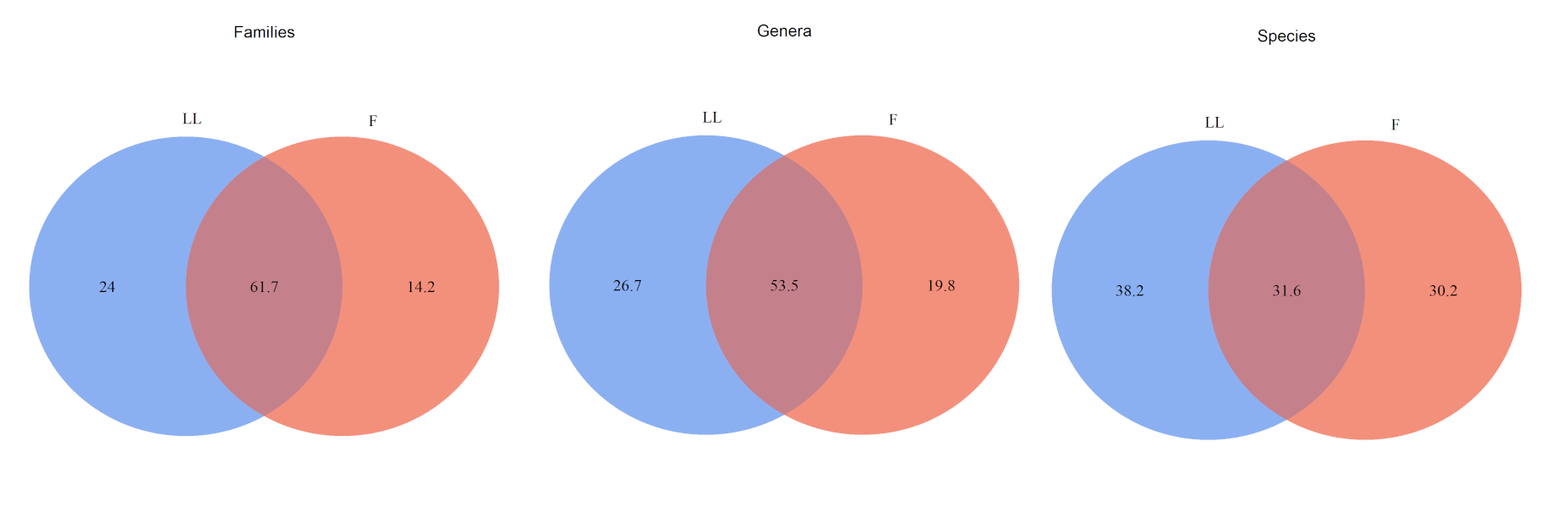

**Figure S3. Venn Diagram for the observed Families, Genera and Species for matched samples.**

**“Gut microbiota diversity analysis.”**

**Figure S4**

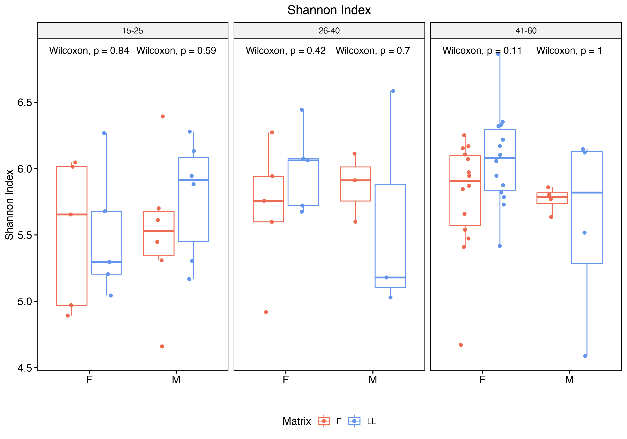

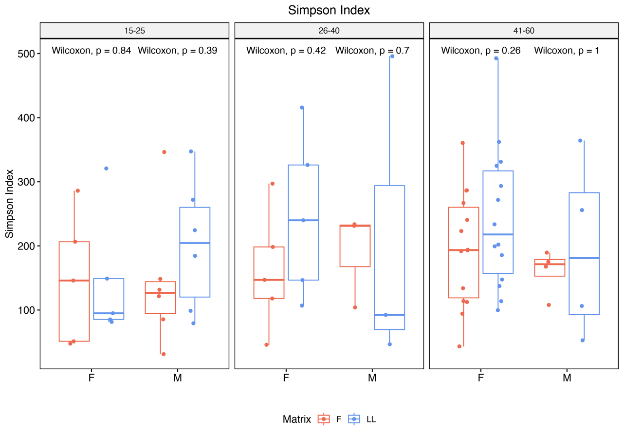

**Figure S4. Alpha diversity comparison between faeces (F) and colonic lavage liquids (LL).** The biodiversity of microbiome was measured using **A)** Shannon diversity index and **B)** Inverse Simpson’s evenness on Observed ASVs following data stratification according to gender and age. Alpha diversity scores were calculated by using normalized data by rarefaction to 68,000 sequences. Each point represents the diversity score for a patient sample. Between-group variations were measured using the Student’s test (S) and the Wilcoxon tests (W) and P values for each alpha diversity measured was reported for each group.

**Figure S5**

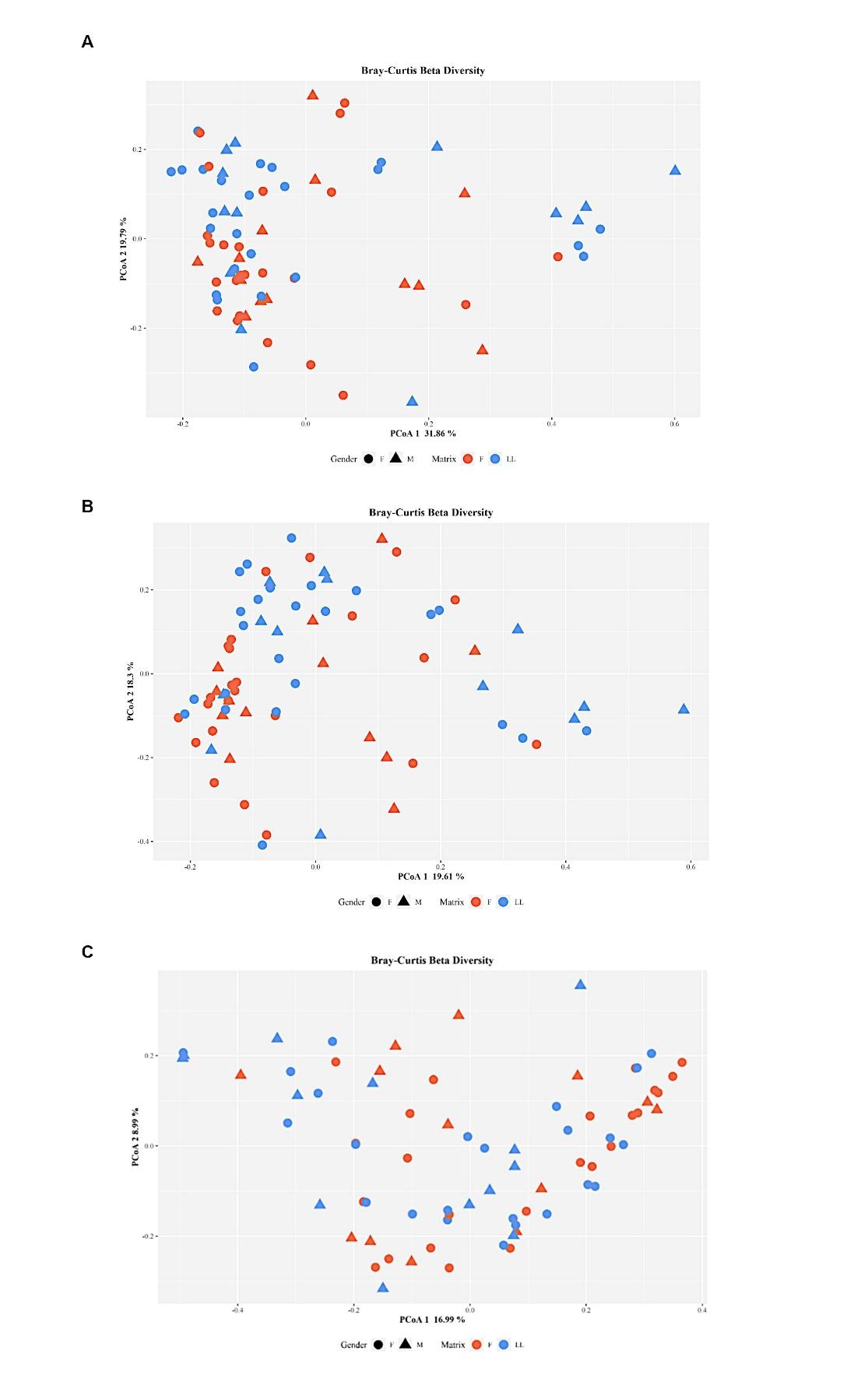

**Figure S5. PCoA plot based on Bray-Curtis dissimilarity matrix at Families (A), Genus (B) and Species levels (C).**

**“Microbial taxa and metabolic pathways correlation.”**

**Figure S6**

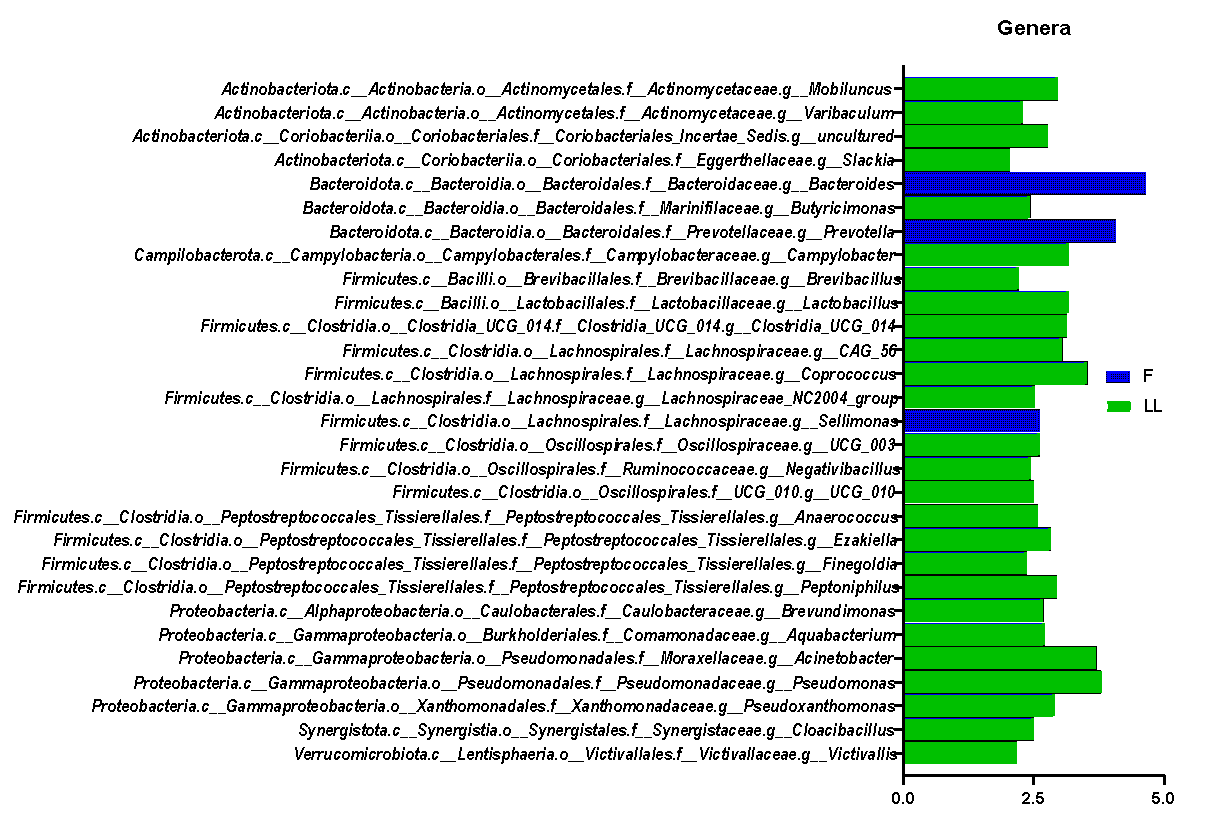

**Figure S6. LEfSe results for Genera.** Genera differently associated with Fecal (blue) and LL (green) samples are shown.

**Figure S7**

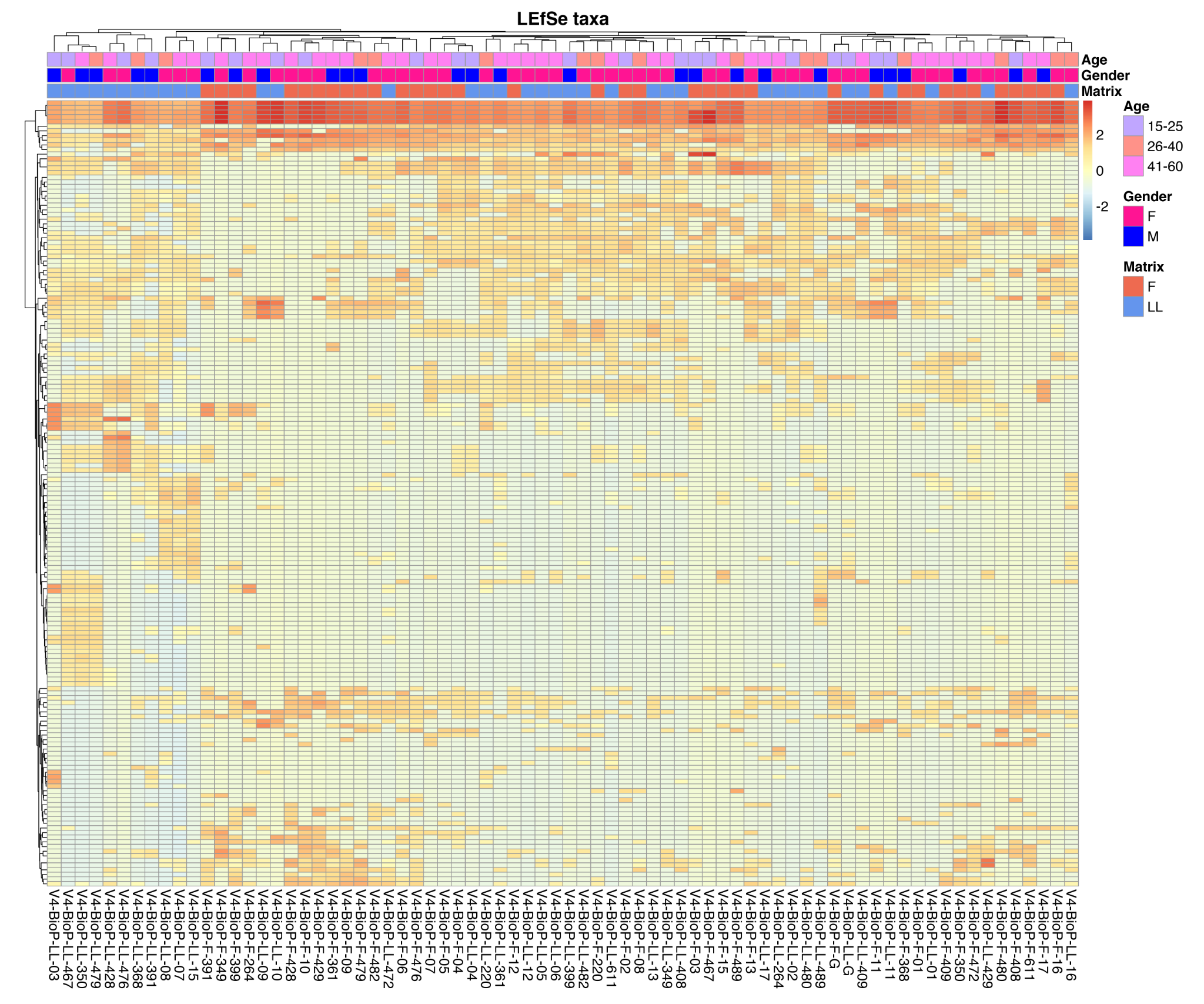

**Figure S7. Heat map of the LEfSe retained taxa.**
